# Socioecological drivers of injuries in female and male rhesus macaques (*Macaca mulatta*)

**DOI:** 10.1101/2023.10.20.563310

**Authors:** Melissa A. Pavez-Fox, Erin R. Siracusa, Samuel Ellis, Clare M. Kimock, Nahiri Rivera-Barreto, Josue E. Negron-Del Valle, Daniel Phillips, Angelina Ruiz-Lambides, Noah Snyder-Mackler, James P. Higham, Lauren J.N. Brent, Delphine De Moor

## Abstract

Competition over access to resources, such as food and mates, is believed to be one of the major costs associated with group living. Two socioecological factors suggested to predict the intensity of competition are group size and the relative abundance of sexually active individuals. However, empirical evidence linking these factors to injuries and survival costs is scarce. Here, we leveraged 10 years of data from free-ranging rhesus macaques where injuries inflicted by conspecifics are associated with a high mortality risk. We tested if group size and adult sex ratio predicted the occurrence of injuries and used data on physical aggression to contextualise these results. We found that males were less likely to be injured when living in larger groups, potentially due to advantages in intergroup encounters. Females, instead, had higher injury risk when living in larger groups but this was not explained by within-group aggression among females. Further, male-biased sex ratios predicted a weak increase in injury risk in females and were positively related to male-female aggression, indicating that male coercion during mating competition may be a cause of injuries in females. Overall, our results provide insights into sex differences in the fitness-related costs of competition and empirical evidence for long-standing predictions on the evolution of group living.

## Introduction

Competition over access to resources is believed to be an important selective pressure for the evolution of group living. By forming groups, animals can gain advantages such as higher success at locating food, more and easily accessible mating opportunities, decreased predation risk and cooperative defence of resources (Jarvis et al., 1998; Ratcliffe and Ter Hofstede, 2005; Silk, 2007; Van Schaik and Van Hooff, 1983). However, life in groups can also be associated with major costs for individuals as a result of competition with conspecifics when valuable resources-such as food and mates-are limited (Terborgh and Janson, 1986; Van Schaik and Van Hooff, 1983; Janson and Goldsmith, 1995). Intense competition in the form of physical aggression can have substantial health costs for individuals because their risk of injury increases (Vogel et al., 2007; Feder et al., 2019). Injuries may indirectly impact reproductive success as animals may need to divert energetic resources to healing (Archie et al., 2014), and can directly impact survival in the case of fatal aggression (Pavez-Fox et al., 2022; Chilvers et al., 2005). Given the fitness costs of injuries, animals are expected to refrain from engaging in physical aggression unless necessary when resources are limited or very valuable (Hammerstein, 1981). Two aspects of a group have been hypothesised to drive the intensity and costs of competition: group size and the operational sex ratio.

Group size might determine the intensity of competition within and between groups. Larger groups have more individuals who need access to the food that is available and so usually suffer from higher levels of within-group competition compared to individuals in smaller groups (Heesen et al., 2014; Balasubramaniam et al., 2014; Gillespie and Chapman, 2001; Blumstein et al., 1999; Marino, 2010). However, when feeding areas can be monopolised and are extensive enough to sustain entire groups, larger groups have a numerical advantage, which can be beneficial for the collective defence of such resources (Cheney and Seyfarth, 1987; McComb et al., 1994). For instance, studies in several species have shown that larger groups are more likely to win between-group encounters than smaller groups (Majolo et al., 2020; Balasubramaniam et al., 2014; Willems et al., 2013; Thompson et al., 2017; Dyble et al., 2019). Differences in life history between the sexes mean that the incentive to compete and the costs/benefits of group size may differ between males and females. In mammals, the fitness of females is mainly limited by access to food, while the fitness of males is mainly limited by access to mates (Trivers, 1972). Increased within-group competition over food access in larger groups has been suggested to impact females more than males (Sterck et al., 1997; Koenig, 2002).Males, on the other hand, tend to be more involved in between-group competition (Smith et al., 2022), whereby resident males collectively defend females or the resources females feed on or against immigration attempts and where larger groups provide a competitive advantage (Cowlishaw, 1995; Majolo et al., 2020; Scarry, 2013). Thus, how group size influences the costs of competition is likely to be sex-dependent.

Another factor suggested to drive competition within a group is the relative availability of sexually active males and females (the operational sex ratio). When the operational sex ratio is skewed, theory predicts there will be higher competition amongst the more abundant sex over access to the less abundant sex (Kvarnemo and Ahnesjo, 1996; Clutton-Brock and Parker, 1992; Emlen and Oring, 1977). For instance, in reindeer (*Rangifer tarandus*), female-female competition for males was higher in a group with a female-skewed operational sex ratio than in a group where the sex ratio was balanced (Driscoll et al., 2022). Similarly, in vervet monkeys (*Chlorocebus pygerythrus*), male-male fights were more frequent in groups with male-skewed operational sex ratios (Hemelrijk et al., 2020). However, when the operational sex ratio is too skewed and the costs associated with aggression are too high, a reduction in intrasexual competition in the abundant sex might be favoured and other strategies could arise (Weir et al., 2011; Rankin et al., 2011). Given that in mammals, females’ damaging potential is usually lower than males - particularly in species with sexual dimorphism (*i.e.,* larger body/canine size in males) - one strategy often adopted by males that might reduce costs associated with male-male retaliation is redirecting the aggression towards females (Clutton-Brock and Parker, 1995; Reale et al., 1996; Smit et al., 2022; Davidian et al., 2022). As a consequence, the operational sex ratio might not only determine costs derived from intrasexual competition but also from inter-sexual aggression.

While the drivers of competition in group-living animals have been well established (Van Schaik and Van Hooff, 1983; Clutton-Brock and Huchard, 2013; McComb et al., 1994; Blumstein et al., 1999; Dyble et al., 2019), there is still scarce empirical evidence for how these factors influence the occurrence of physical aggression, with consequences for injuries and fitness. Quantifying the consequences of contest competition and its fitness outcomes has proven difficult in most wild systems where injuries or body damage can be caused by predators and not be the direct result of competition with conspecifics. Obtaining behavioural information from large wild groups and estimating the operational sex ratio when there are roaming or dispersing males can also be challenging (Kappeler, 2017). Further, given the differences in life history between the sexes, the costs and drivers of competition often are considered separately for males and females, even though there is mounting evidence that mating competition can also result in sexual conflict (Davidian et al., 2022; Smit et al., 2022; Baniel et al., 2017).

To quantify the fitness costs of contest competition, here, we explore the socioecological drivers of injuries in free-ranging female and male rhesus macaques living in Cayo Santiago, Puerto Rico. Rhesus macaques live in multi-female multi-male societies where females are philopatric and males disperse at sexual maturation (Thierry et al., 2004). Females form strict despotic dominance hierarchies where rank is maternally inherited (Chikazawa et al., 1979). Males acquire rank via a queuing system where group tenure determines their social status (Manson, 1995; Kimock et al., 2022).Rhesus macaques have a polygynandrous mating system with high synchrony in females’ fertile phases, reducing the monopolisation potential of males (Dubuc et al., 2011). As a consequence, male rhesus often rely more on indirect forms of competition, such as sperm competition, endurance rivalry, sneaky copulations and female coercion (Higham et al., 2011; Higham and Maestripieri, 2014; Manson, 1994), rather than direct male-male conflict (Kimock et al., 2022). There are no predators of rhesus macaques on the Cayo Santiago island, and injuries are primarily caused by conspecifics. Injuries have been linked to a 3-fold decrease in survival probability in this population for both sexes (Pavez-Fox et al., 2022), providing the opportunity to test the fitness-related costs of competition. Demographic information is collected monthly providing accurate information on group membership and sex ratio. Social groups are naturally formed and vary in size from 26 to nearly 300 adults. Although the population is food provisioned, competition over monopolizable food and water stations frequently occurs, where high-ranking macaques spend on average more time feeding and drinking than low-ranking animals (Balasubramaniam et al., 2014).

To determine the socioecological drivers of injuries in this population, we tested for sex-specific effects of group size and adult sex ratio (sex ratio henceforth), a proxy of operational sex ratio, on the occurrence of injuries. Because injuries were collected opportunistically, we do not have information on who caused the injury, although due to the lack of predators on the island we can be confident that all injuries were inflicted by conspecifics. Given this, we used long-term behavioural observations of physical aggression, where we have data on the sex of both the victim and aggressor, to contextualise the injury data. Specifically, we looked at how sex ratio and group size predicted intrasexual and intersexual physical aggression to help infer the cause of injuries and therefore the underlying drivers of competition in this system. We predicted that in larger groups the risk of injury (*i.e.,* probability of being injured) would be higher for females and lower for males. We predicted that females would experience higher injury risk in larger groups as a result of higher within-group female-female (FF) feeding competition (Wrangham et al., 1993; Chapman and Chapman, 2000), thus we further tested if FF physical aggression (*i.e.,* probability of being physically aggressed) was higher in larger groups. For males, we expected that those living in larger groups would have reduced injury risk because having a numerical advantage translates into better chances of winning between-group encounters (Koenig et al., 2013; Janson and Goldsmith, 1995). Given that the behavioural data included only within-group interactions, we could not test patterns of between-group aggression. Instead, we tested whether male-male (MM) physical aggression was influenced by group size to rule out within-group MM competition as a driver of injuries. For sex ratio, if classic socioecological predictions apply to mating competition in rhesus macaques, we expected that in groups with skewed sex ratios, those individuals of the sex in minority would have higher injury risk due to more intense competition over mates (Kvarnemo and Ahnesjo, 1996). That is, male injury risk and MM physical aggression might increase as the sex ratio becomes more male-biased while female injury risk and FF physical aggression are expected to increase as the sex ratio becomes more female-biased. However, we do not necessarily expect these classic predictions for sex ratio in rhesus macaques, because males often rely on indirect forms of competition (Higham et al., 2011; Higham and Maestripieri, 2014; Kimock et al., 2022) and because females live in philopatric societies where the incentive to compete aggressively over mates against their kin is typically low (Davidian et al., 2022).As such, we have alternative rhesus-specific predictions. We expected that the local availability of mating partners would not determine injury risk in males, because males are not expected to frequently engage in direct MM competition over females and there is therefore no reason to expect an effect of sex ratio on MM physical aggression. Instead, we expected that females would have higher injury risk when females are more scarce (male-biased sex ratio). This is because rhesus macaques are sexually dimorphic and male coercion has been reported (Manson, 1994). This means that male-female (MF) physical aggression would be expected to be higher in groups with a male-biased sex ratio where males direct their aggressive behaviours towards females when competition over females increases. Finally, we predicted that FF physical aggression would not be influenced by sex ratio as female rhesus macaques are not expected to fiercely compete over access to males against their kin. All predictions are laid out in Table 1.

**Table 1:** Predictions for the socioecological drivers of injuries in rhesus macaques.

| Sex | Group size | Sex ratio classical | Sex ratio rhesus |
| --- | --- | --- | --- |
| Females | ↑ injury risk in larger groups | ↑ injury risk when female-biased | ↑ injury risk when male-biased |
|  | ↑ FF aggression in larger groups | ↑ FF aggression when female-biased | ↑ MF aggression when male-biased, No effect on FF aggression |
| Males | ↓ injury risk in larger groups | ↑ injury risk when male-biased | No effect on injury risk |
|  | No effect on MM aggression within groups | ↑ MM aggression when male-biased | No effect on MM aggression within groups |

Sex ratio classical: general predictions of socioecological models, Sex ratio rhesus: predictions specific to our study system, group size: number of individuals in the group (4 years and older), sex ratio: number of adult females per male during the mating season, aggression: physical aggression, ‘FF’: female-female, ‘MM’: male-male, ‘MF’: male-female.

## Methods

### Study subjects

Study subjects were free-ranging male and female rhesus macaques living on Cayo Santiago island, Puerto Rico. The island is home to a population of *∼* 1800 individuals living in 6 to 12 mixed-sex naturally formed social groups. The Cayo Santiago field station is managed by the Caribbean Primate Research Center (CPRC), which monitors the population daily and maintains the long-term (*>*75 years) demographic database including data on births, deaths and social group membership for all animals (Kessler and Rawlins, 2016).Macaques are individually identified based on tattoos located on their chest and legs. Animals have *ad-libitum* access to food and water, the island is predator-free and there is no regular medical intervention for sick or wounded individuals. Here we included data on male and female macaques that were alive between 2010 and 2020. We focused on individuals aged 4 years and above (age range: 4 - 28 years), as animals of both sexes have typically reached sexual maturity at that age (Zehr et al., 2005; Bercovitch and Berard, 1993). We restricted our sample to animals belonging to social groups for which we had data on injury occurrence and agonistic behavioural observations (*n* = 6 social groups). The groups analysed varied in size from 26 to 288 animals and sex ratios (*n* females/ *n* males) ranged from 0.5 to 4.5 (Fig. S1).

### Observation of Injuries

Since 2010, the CPRC staff have been collecting opportunistic observations on the incidence and recovery from injuries during the daily monitoring of social groups for demographic purposes. Data collection was carried out mainly by the veterinary technician complemented by information from other experienced staff during the working hours of the field station (7:30 to 14:00 from Monday to Friday). If an individual was observed to be wounded or displaying signs of injury (*e.g.,* limping) the staff member recorded the individual ID and if the injury was visible, the type of injury (*e.g.,* puncture, scratch), the area of the body affected, whether the injury was recent or old based on the presence of scars, and if possible, an estimate of the wound size. Records for each individual were updated every time the observers encountered the wounded individuals during the daily census. Here we included all records for visible injuries including bites, scratches, abrasions and cuts along with other more severe injuries such as exposed organs and fractures. We decided to exclude cases where injuries could be inferred but not observed, such as limping or abscesses as these could also be caused by infection unrelated to injury. We excluded injury records from two full years (2015 and 2016), a period for which the veterinary technician was not regularly at the field site, which may have led to biases in the few groups sampled during those years. Our sample consisted of 908 injuries collected from September 2010 to April 2020 on 521 unique individuals (*n* females = 267, *n* males = 254).

### Collection of aggression data

We collected behavioural data using focal samples based on a previously established ethogram (Brent et al., 2014) from twenty different group years (group F 2010-2017, group HH 2014 and 2016, group KK 2015 and 2017, group R 2015 and 2016, group S 2011 and 2019, group V 2015-2019). Across the 10 years of study, two external events in 2018 and 2020 - Hurricane Maria and the COVID-19 pandemic, respectively - precluded the collection of focal data. These years were excluded from the aggression analyses. In total, this resulted in 748 adult individuals (422 females and 326 males) whose ages ranged between 4-28 years old (mean = 10.7). Behavioural data were collected using 10-min (17 group years) or 5-min (3 group years) focal animal samples between 07:30 and 14:00. We stratified sampling to ensure balanced data collection on individuals throughout the day and over the year. During focal sampling, dyadic agonistic encounters where the focal animal was involved were recorded, along with the identity of the aggressor and victim. We recorded all agonistic interactions, including submissions, threats, non-contact aggression (*e.g.,* charge, chase), and physical aggression (*e.g.,* bite, hit). Given that the purpose of our study was to use the aggression data to contextualise the occurrence of injuries, we considered only data on physical aggression, which is more likely to lead to an injury. From January 2010 to October 2019, we recorded 18880 aggression events including 522 physical aggression.

### Quantifying injury risk and aggression rates

The injury dataset included the 521 animals that were recorded injured in addition to 1001 uninjured animals (*n* uninjured females = 525, *n* uninjured males = 476). Uninjured individuals consisted of all sexually mature individuals who were alive during the period of study, *i.e.,* between 2010 and 2020 excluding 2015 and 2016 to match data on injured animals. Given that the average recovery time for an injury was 43 days and the average time elapsed between consecutive injury records was 42.17 days, the dataset was formatted in a way that each row represented a two-month interval period (*i.e.,* bimonthly interval). By formatting the data this way we could be confident that injury records occurring in different rows were more likely to be independent (for details see SI: Pavez-Fox et al. 2022). An individual’s injury status during each bimonthly interval they were alive was coded as a binary variable where 1 = injured and 0 = uninjured.

The aggression dataset included the 748 male and female macaques for which focal data were collected. Given that our questions were sex-specific, we split this dataset by the sex of the focal animal resulting in 438 physical aggression events in a total of 422 females and 84 physical aggression events in a total of 326 males. We focused specifically on physical aggression received by the focal animal. Each row represented a bimonthly interval to match the format of the injury data. Given that an individual rarely received physical aggression more than once in a given bimonthly interval (Fig. S2), we coded an individual’s aggression status as binary, where 1 = physically aggressed and 0 = not physically aggressed. Depending on the question, we split this dataset based on the sex of the victim and the aggressor.

### Statistical analyses

We ran all the models in a Bayesian framework using the brms R Package (Burkner, 2021).Therefore, evidence of an effect was determined based on the degree of overlap between the credible interval (CI) and zero (*i.e.,* 89% non-overlapping reflecting strong evidence of an effect). Given that all the dependent variables were coded as binary, models were fit using a Bernoulli distribution. All continuous predictors were z-scored. In all the models we included random intercepts for individual ID to account for repeated measures and for the specific bimonthly interval within the study period. We assumed normal distributions for priors (mean = 0, SD = 1) and ran 10000 iterations in all the models. Model assumptions and posterior predictive checks were done using the ‘ppcheck’ built-in function from the brms package. Marginal effects were calculated using the emmeans R package(Lenth et al., 2018). We reported means as point estimates, standard error (SE) and 89% credible intervals of the posterior distribution. For marginal effects, we reported the median and the 89% highest posterior density interval (HPD).

### Group size and sex as drivers of injuries

#### Effect of group size on injury risk

To test whether group size predicted the probability of an individual being injured, we built a model where the dependent variable was an individual’s injury status (1/0) and the independent variables included group size, the individual’s sex and the reproductive season (1 = mating, 0 = non-mating) in a given bimonthly interval. Because our predictions were sex-specific, we included an interaction term between group size and sex. Using demographic records, we computed group size as the number of adults (4 years and above) that were alive in a subject’s group in a given bimonthly interval. We specifically determined a group’s size at the middle of the interval (end of the first month), thus if an individual reached 4 years of age or died during the second month, this was only reflected in the following bimonthly interval. We determined the reproductive season following (Hoffman et al., 2008). Briefly, we first computed the mean birth date ± 2 SD for each year. The start of the birth season was defined as the first birth date and the end as the last birth date. The beginning of the mating season was determined by subtracting the gestation period of rhesus macaques (165 days; Silk et al. 1993) from the start of the birth season, and the end of the mating season was determined by subtracting the gestation period from the end of the birth season. If the middle of the bimonthly interval fell outside the mating season it was considered part of the non-mating period.

#### Effect of group size on female-female aggression

To test whether FF competition might be a driver of injuries in females living in larger groups, we focused on female-female aggression data. The dependent variable was female aggression status (0/1) and we included the number of females in the group, the reproductive season and an offset term for sampling effort (*i.e.,* hours an individual was focal-followed) as independent variables in the model. We used the number of females in the group rather than group size as a predictor in the model because the former better reflects FF competition and these two metrics were strongly correlated (Fig. S3A, Pearson’s *R* = 0.94, p *<* 0.01). Using the same model specifications, we additionally tested whether group size predicted MF physical aggression (where the victims were females and the aggressors were males) to rule out other drivers of injuries in females related to within-group competition.

#### Effect of group size on male-male aggression

To test our prediction that within-group MM competition was not a driver of injuries in smaller groups we focused on male-male aggression data. We tested if the number of males in a group, which was positively correlated to group size (Fig. S3B, Pearson’s *R* = 0.97, p *<* 0.01),predicted a male’s risk of physical aggression from other males in his group. The dependent variable was a male’s aggression status (0/1) and the independent variables were the number of males in a group, the reproductive season and an offset term for sampling effort. The occurrence of FM physical aggression was very rare (only 9 cases across the 10 years), thus we disregarded within-group female aggression as a driver of injuries in males.

### Sex ratio and sex as drivers of injuries

#### Effect of sex ratio on injury risk

To test whether the sex ratio predicted the probability of an individual being injured we built a model where the dependent variable was an individual’s injury status (0/1) and included sex ratio and sex as independent variables. Given that our predictions were sex-specific, we included an interaction term between the sex ratio and sex. We computed the sex ratio as the number of females (4 years and above) per male in the subject’s group in a given bimonthly interval. Therefore, smaller numbers would indicate a male-biased sex ratio while larger numbers would indicate a female-biased sex ratio. For these analyses, we focused on the mating season, to have a better estimate of sexually active individuals and to make sure that the socioecological driver of injuries was competition for mates. As with group size, we determined a group’s sex ratio at the middle of the bimonthly interval, thus if an individual reached 4 years of age or died during the second month, this was only reflected in the following interval.

#### Effect of sex ratio on male-female aggression

To test if MF coercion was a driver of injuries in females we focused on aggression data where the victims were females and the aggressors were males. As a dependent variable, we included a female’s aggression status (0/1) and as independent variables, the sex ratio and an offset term for sampling effort. As above, we focused on the mating season for this analysis to make sure that mating competition was the driver of physical aggression.

*Effect of sex ratio on female-female aggression*. To test whether FF competition over males was a driver of injuries in females we focused on data where the aggressor and the victim were females. As above, we restricted this analysis to the mating season. The dependent variable was a female’s aggression status (0/1) and independent variables included sex ratio and an offset term for sampling effort.

#### Effect of sex ratio on male-male aggression

To test if MM competition over females was a driver of injuries in males we focused on male-male aggression data during the mating season. The dependent variable was a male’s aggression status (0/1) and predictors included sex ratio and an offset term for sampling effort. To rule out the possibility that young and old females might not be attractive partners for males to compete over (as we consider all females over 4 years of age), we also tested the effect of the adult sex ratio considering only the number of prime-age females (6-17 years; Lee et al. 2021) per male in the group.

## Results

### Group size and sex as drivers of injuries

#### Effect of group size on injury risk

In support of our predictions, we found a sex-dependent effect of group size on injury risk (Fig. 1A; Log-Odds group size*sexM = −0.36, SE = 0.08, 89% CI = −0.49, −0.23; Table S1). Females were 53% more likely to be injured for every one SD (*∼* 59 individuals) of increase in group size (marginal effect: Log-Odds females = 0.14, 89% HPD = 0.025, 0.26). In the case of males, an increase in one SD in group size was associated with a reduction of 44% in the probability of being injured (marginal effect: Log-Odds males = −0.22, 89% HPD = −0.33, −0.11) (Fig. 1B).

**Figure 1:**
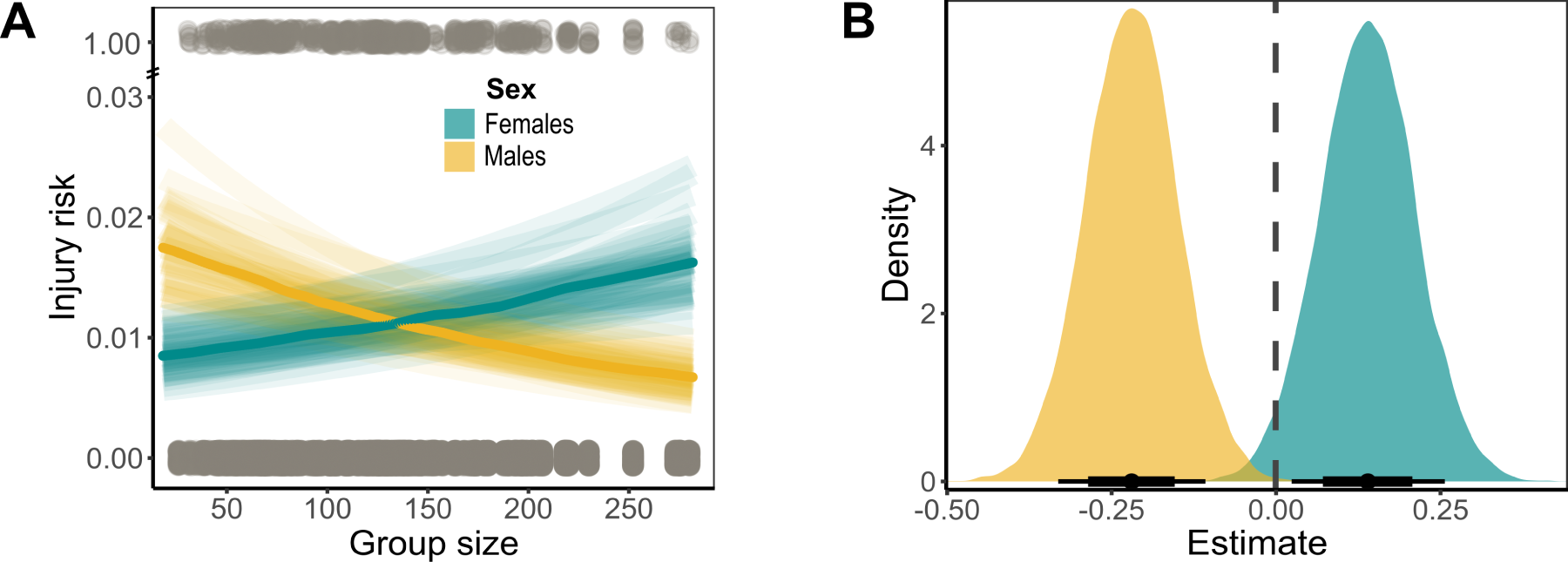
Sex-dependent effect of group size on injury risk. **A)** Predicted values of injury risk for females (cyan) and males (yellow) as a function of group size. The darker line indicates the median and the lighter lines show the 89% CI. Raw data are depicted with grey points (top: injured, bottom: uninjured). **B)** Posterior distributions for marginal effects of group size on male and female injury risk. Whiskers indicate the median, 89% CI (thinner line) and 66% CI (thicker line).

#### Effect of group size on female-female aggression

Contrary to our prediction, females living in groups with more females (*i.e.,* larger groups) were not more likely to be physically aggressed by other females in the group (Fig. 2 top panel; Log-Odds fem count = −0.09, SE = 0.08, 89% CI = −0.22, 0.03; Table S2). We interpret this to mean that there is no evidence of FF competition driving injuries in larger groups. We could also rule out MF physical aggression with group size, as females were less likely to be physically aggressed by males as group size increased (Fig. 2 middle panel; Log-Odds group size = −0.14, SE = 0.08, 89% CI = −0.26, −0.01; Table S3).

**Figure 2:**
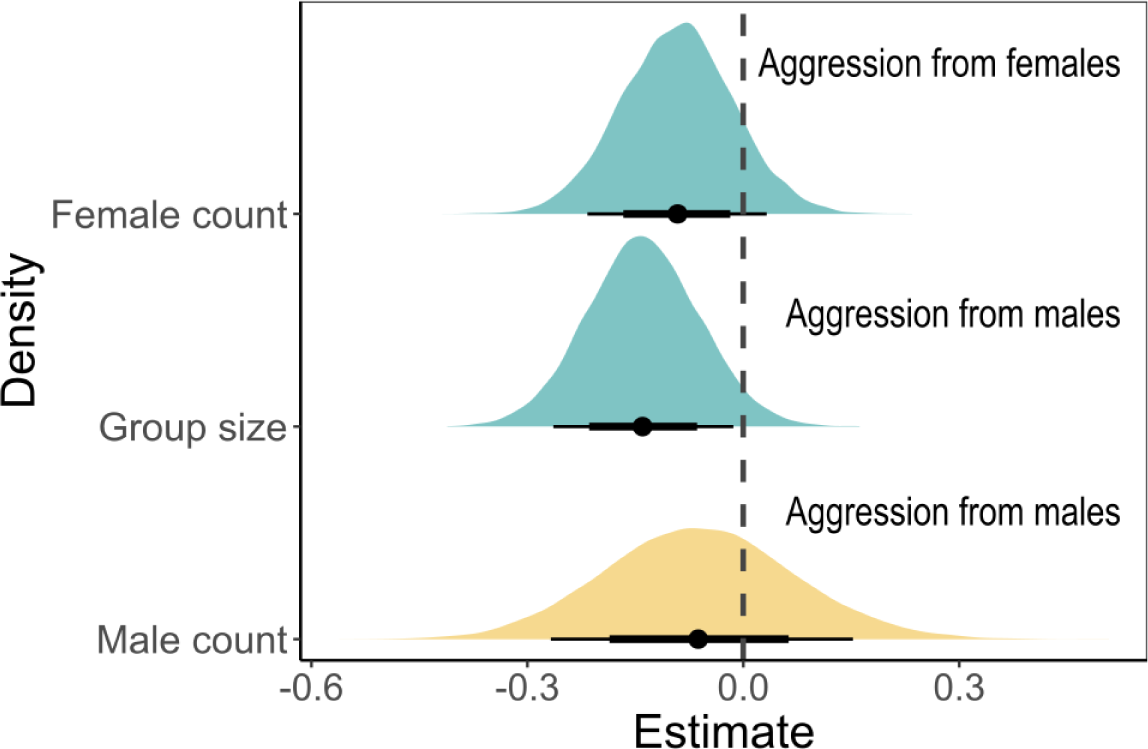
Sex-specific drivers of injuries with group size. **A)** Posterior distributions of estimates from models testing the effect of the number of females in a group on FF physical aggression (top panel), group size on MF physical aggression (middle panel), and the number of males on MM physical aggression (bottom panel). Female victims are depicted with cyan and male victims with yellow. Whisker indicates the median, 89% CI (thinner line) and 66% CI (thicker line).

#### Effect of group size on male-male aggression

As predicted, we did not find evidence of an effect of group size on MM physical aggression within groups. The number of males in a group did not predict the likelihood of a male receiving physical aggression from other resident males (Fig. 2 bottom panel; Log-Odds male count = −0.06, SE = 0.13, 89% CI = −0.27, 0.15, Table S4).

### Sex ratio and sex as drivers of injuries

#### Effect of sex ratio on injury risk

We found a sex-dependent effect of sex ratio on an individual’s injury risk (Fig. 3A; Log-Odds sex ratio*sexM = 0.17, SE = 0.08, 89% CI = 0.04, 0.3; Table S5). Contrary to our rhesus-specific and classical predictions, males who lived in groups with female-biased sex ratios were more likely to be injured. For every increase in one SD of sex ratio (*∼* 0.5 increase in females relative to males), males experienced a 53% increase in their likelihood of being injured (marginal effect: Log-Odds males = 0.12, 89% HPD = 0.01, 0.21). Females were more likely to be injured when living in groups with a male-biased sex ratio, but this relationship was weak as the credible interval overlapped with zero (marginal effect: Log-Odds females = −0.05, 89% HPD = −0.16, 0.06) (Fig. 3B).

**Figure 3:**
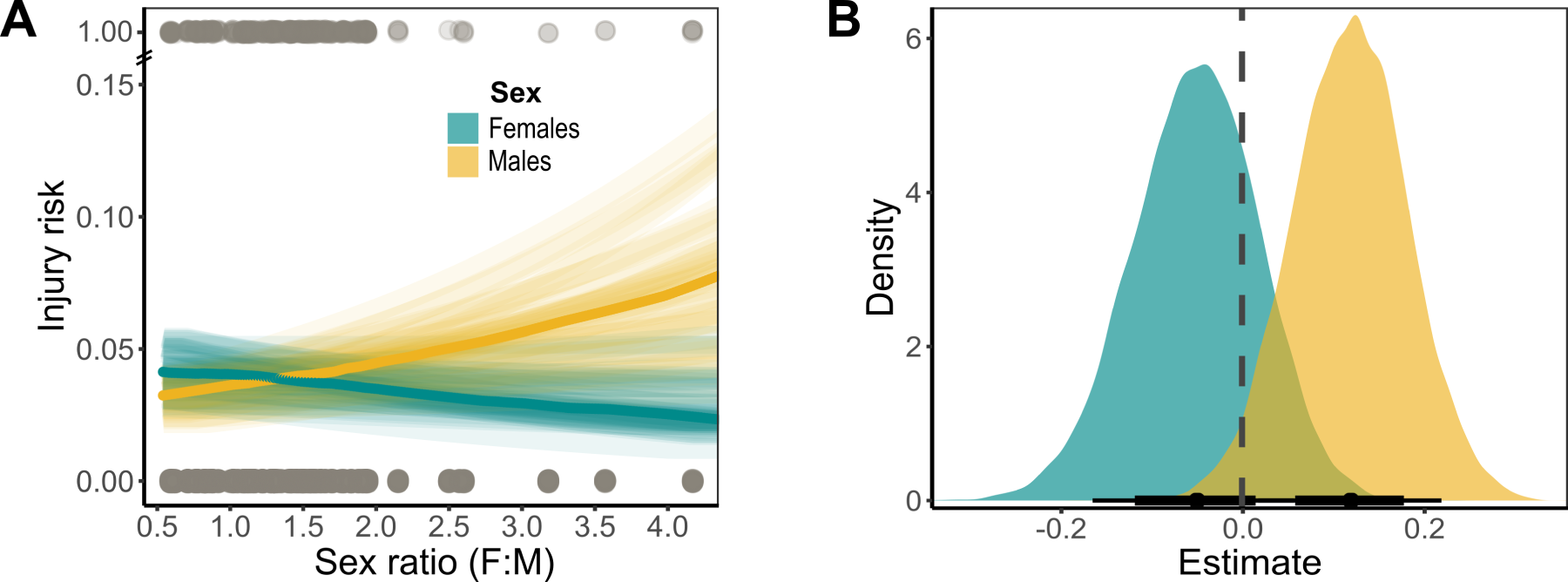
Sex-dependent effect of adult sex ratio on injury risk. **A)**Predicted values of injury risk for females (cyan) and males (yellow) as a function of adult sex ratio (*i.e.,* number of females per male during the mating season). The darker line indicates the median and the lighter lines show the 89% CI. Raw data are depicted with grey points (top: injured, bottom: uninjured). **B)** Posterior distributions for the estimates of adult sex ratio on male and female injury risk. Whiskers indicate the median, 89% CI (thinner line) and 66% CI (thicker line).

#### Effect of sex-ratio on male-male aggression

We did not find evidence for MM competition over females driving injuries, as males were not more likely to receive physical aggression by resident males when living in groups with a male-biased operational sex ratio (Fig. 4A top panel; Log-Odds sex ratio = 0.1, SE = 0.17, 89% CI = −0.19, 0.37, Table S6). This result holds even when only prime-aged females were considered in the computation of sex ratio (Log-Odds sex ratio = 0.04, SE = 0.19, 89% CI = −0.27, 0.34, Table S7).

**Figure 4:**
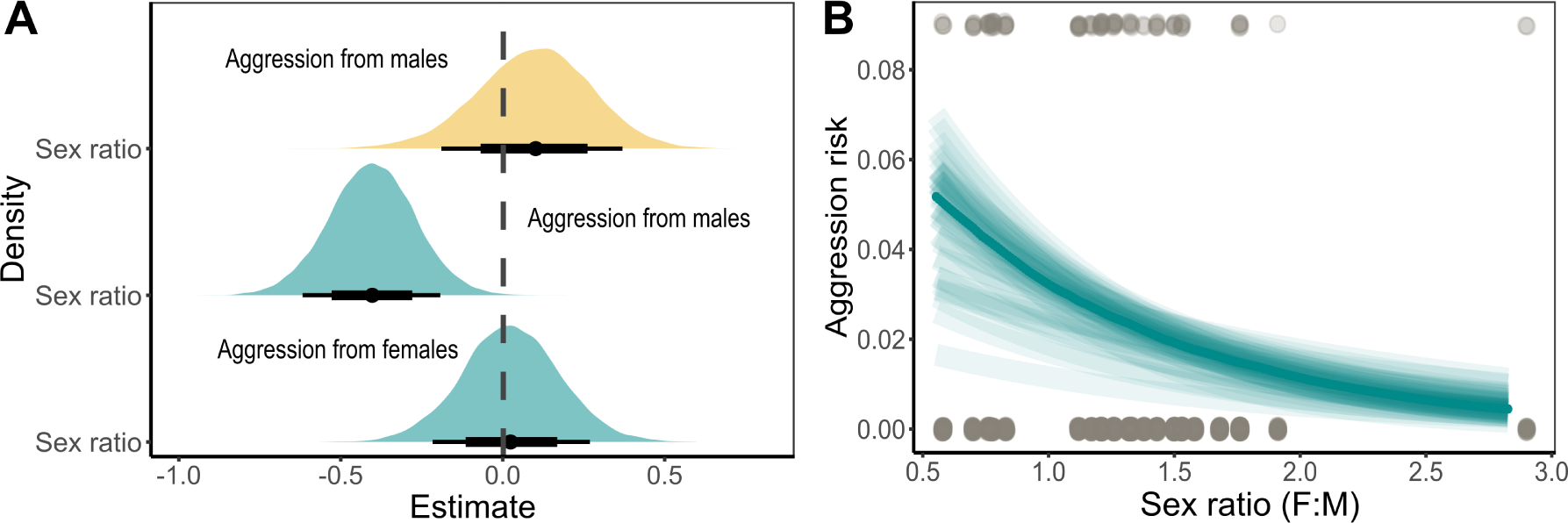
Sex-specific drivers of injuries with sex ratio. **A)** Posterior distributions of estimates from models testing the effect of sex ratio (number of females to males) on MM physical aggression (top panel), sex ratio on MF physical aggression (middle panel), and sex ratio on FF physical aggression (bottom panel). Female victims are depicted with cyan and male victims with yellow. Whisker indicates the median, 89% CI (thinner line) and 66% CI (thicker line). **B)** Predicted values for the risk of physical aggression from males to females as a function of sex ratio. The darker line indicates the median and the lighter lines show the 89% CI. Raw data are depicted with grey points (top: physically aggressed, bottom: not physically aggressed).

#### Effect of sex ratio on male-female aggression

Consistent with our rhesus-specific prediction, male-to-female physical aggression was negatively associated with the relative availability of females in a group. For every one SD decrease in sex ratio (*∼* 0.5 decrease in the number of females relative to males), females were 40% more likely to be physically aggressed by males (Fig. 4A middle panel, Fig. 4B; Log-Odds sex ratio = −0.4, SE = 0.13, 89% CI = −0.62, −0.19, Table S8).

#### Effect of sex ratio on female-female aggression

We found no evidence of FF competition for males driving injuries. As predicted for rhesus macaques, during the mating season females were not more likely to be physically aggressed by other females in groups when the relative availability of males was low (*i.e.,* female-biased sex ratio) (Fig. 4A bottom panel; Log-Odds sex ratio = 0.02, SE = 0.15, 89% CI = −0.22, 0.27, Table S9).

## Discussion

In this study, we tested predictions derived from socioecological theory on the sex-specific drivers of injuries. As predicted, we found that living in larger groups may confer a competitive advantage to males but a cost to females. Males living in larger groups were less likely to be injured compared to males in smaller groups, whereas females had a higher risk of injury in larger groups. Further, we found that female aggression was not a driver of female injury in this population but instead, our results pointed to the role of male coercion during mating competition. In males, we found no evidence of injuries being driven by within-group aggression, suggesting that injuries were likely caused during inter-group encounters.

Taken together these results provide rare evidence of fitness-related costs associated with classic predictors of socioecological models.

### How does group size impact injury risk?

As predicted by socioecological models and life-history traits (Koenig, 2002; Scarry, 2013; Trivers, 1972), we found clear sex differences in how group size predicts injury risk. We discuss these results and the possible socioecological drivers for each sex separately below. Females living in larger groups had a higher risk of injury than females in smaller groups.

However, contrary to our prediction, we found no evidence that this was driven by within-group FF competition, as females in larger groups did not receive more physical aggression from other female group members. Larger groups are believed to impose major energetic constraints, particularly for females, which require high food intake to fulfil the costs of pregnancy and lactation (Markham and Gesquiere, 2017; Trivers, 1972).As a consequence, females are expected to compete more intensely for food when living in larger groups (Sterck et al., 1997; Koenig, 2002). Yet, our results suggest that this might not be the case in female rhesus macaques at the Cayo Santiago field station. This could be because animals in this population are food-provisioned, thus feeding resources might not be as limited or restricted as in wild populations, reducing the incentive for high-cost physical aggression. However, given that females do engage in conflict over food in this population (Balasubramaniam et al., 2014),a complementary, or possibly even alternative explanation, is that the despotic dominance hierarchy that characterises females of this species mediates access to resources and reduces the need for physical aggression (Thierry et al., 2004; Holekamp and Strauss, 2016). In support of the idea that elevated competition in larger groups might be more apparent through non-physical aggression, we found - in a supplementary analysis - that females living in larger groups were more likely to receive non-physical aggression by other females compared to females living in smaller groups (Fig. S4, details in SI).

But if not female-female aggression, what is the source of injuries for females living in larger groups? One possible explanation is male aggression. However, we found the opposite pattern as MF physical aggression decreased with group size. Evidence of reduced male aggression toward females in larger groups could be a consequence of the fact that females in larger groups tend to have more kin and therefore more support against males in agonistic encounters. Together these results show that group size could not explain within-group physical aggression patterns that match higher injury risk in females living in larger groups. This, in turn, suggests that injuries for females in these groups might be the result of intergroup aggression. Some studies in primates have shown that females may participate in intergroup coalitionary aggression more than males (Martınez-Iñigo et al., 2021) and that they are also more likely to engage in intergroup conflict when they have more support from male group members Arseneau-Robar et al. (2017). Further investigation is required to determine the incentives for participation in intergroup aggression in female rhesus macaques.

Males had a lower injury risk when living in larger groups. Given that the number of males in a group did not predict the risk of physical aggression between resident males, these results suggest that the source of injuries likely comes from intergroup encounters. In line with our predictions and results from previous meta-analyses in mammals where the number of males was associated with the resource-holding potential of a group (Smith et al., 2022; Majolo et al., 2020),our results provide indirect evidence that larger groups might confer a collective competitive advantage to males. Males from many mammal species have been shown to engage more often than females in intergroup encounters, possibly as a strategy to keep other males away from female group members (Jordan et al., 2007; Cooksey et al., 2020), or to defend the feeding resources (Fashing, 2001; Furrer et al., 2011; Scarry, 2013). Whether the cost of living in smaller groups comes from injuries during collective encounters between groups or during male immigration attempts, where more males might be better able to deter immigration without physical aggression remains an open question.

### How does sex ratio impact injury risk?

Contrary to classic predictions of theoretical models where skewed sex ratios are proposed to lead to fierce intrasexual mating competition (Kvarnemo and Ahnesjo, 1996), and also to our rhesus-specific predictions (Table 1), we found that males had higher injury risk when the relative availability of females was higher (*i.e.,* female-biased sex ratio). We also found weak evidence for an effect of sex ratio on female injury risk. As above, we discuss these results and the possible socioecological drivers in a sex-specific manner.

We found that in groups where males outnumber females, competition among males was not associated with injury risk or heightened physical aggression during the mating season. These results support our rhesus-specific predictions and previous research suggesting that despite moderate levels of sexual dimorphism, contest competition for mates between resident male rhesus macaques is not common (Higham and Maestripieri, 2014; Kimock et al., 2022). Instead, rhesus macaque males are believed to rely on strategies of indirect competition, such as sperm competition, endurance rivalry (Higham et al., 2011), group tenure (Manson, 1995), sneak copulations (Higham and Maestripieri, 2014), and to a lesser extent, female coercion and mate-guarding (Manson, 1994).However, contrary to our predictions, we found that males were more likely to be injured in groups with a female-biased sex ratio. Males in these groups may be more likely to suffer injuries if the higher relative abundance of females makes the group more attractive to immigrant and outsider males, especially if there are fewer males to resist immigration attempts (Alberts and Altmann, 1995). Indeed, males in this population usually disperse during the reproductive season (Hoffman et al., 2008) and may incur higher costs of injuries when doing so (Kimock et al. *in prep.*).

We found no evidence that female mating competition might result in injuries. Consistent with our rhesus-specific predictions but contrary to classical socioecological theory, we found that sex ratio did not predict physical aggression among females. As highlighted by Davidian et al. (2022),there might be strong selective pressures for reduced intrasexual mating competition in most female mammals. The incentive to physically compete over males may be low as sharing mating opportunities with other females is not as costly as it is for males (although there might be some cases where female-female mating competition does occur; Baniel et al. 2018). Female philopatry may favour the use of less costly means of competition to reduce physical aggression against kin (Young and Bennett, 2013). In line with this, we found in a supplementary analysis that as the group becomes more female-biased, and thus FF mating competition is expected to be higher, non-contact aggression among females increases (Fig. S5, Table S11; details in Supplementary). Further, physical aggression and its consequences may be too costly for females given their higher energetic demand for reproduction (Trivers, 1972). More specifically for rhesus macaques, female extra-group copulation (Manson, 1992) and low risk of infanticide (Camperio, 1984), might further reduce the need to compete fiercely over mating opportunities with resident males (Baniel et al., 2018).

We found some support for male coercion as a possible cause of injuries in females. Females living in groups with a male-biased sex ratio were more likely to be physically aggressed by males (although we did not find evidence for a similar effect on female injury risk). These results together provide partial support for our rhesus-specific prediction and previous evidence suggesting that males of this species and others, may rely on coercive strategies when competition for females is intense (Bercovitch, 1997; Bercovitch et al., 1987; Smit et al., 2022; Baniel et al., 2017).One likely explanation for resident rhesus males relying on coercive strategies is to deter female mate choice, as female rhesus macaques prefer to mate with outsider males, potentially due to benefits derived from increasing genetic variability or quality (Manson, 1992). The lack of evidence for an effect of sex ratio on female injury risk might also be attributed to reduced sample size, as our injury results trended in the expected direction but unlike the analyses exploring injury risk with group size, we considered only injuries that occurred within the mating season, which substantially reduced our sample size. Alternatively, it is also possible that males rely on less severe forms of physical aggression when coercing females in their groups (like slaps or hits), which might not lead to injuries. Although we can not confidently conclude that male physical aggression results in females being injured, our results suggest that aggression from resident males could be one source of injuries in female rhesus macaques.

## Conclusion

In this study, we showed a sex-dependent effect of group size and sex ratio on the occurrence of injuries, which have been shown to have detrimental survival consequences. Our group size results demonstrate that within-group intrasexual competition might not lead to injuries in males or females, suggesting instead that intergroup conflict may play a role in individual injury risk and mortality in this population. Moreover, we also found that male coercion might be one source of female injury during mating competition. While the Cayo Santiago population is food-provisioned and predator-free, which might reduce the need for contest competition over food and mates, the episodes of physical aggression and injuries we detected here suggest that the fitness costs of competition in wild populations might be even higher. Overall, our study provides empirical evidence for fitness-related costs of fundamental aspects of social organisation.

## Acknowledgments

We thank the CPRC for the permission to undertake research on Cayo Santiago, along with Edgar Davila, Julio Resto, Bianca Giura, and Giselle Caraballo, who assisted in the collection of injury data. Additionally, we thank members of the Centre for Research on Animal Behaviour (CRAB) for their helpful suggestions.

## Data and code availability

Data used in the analyses will be made available upon acceptance. R code used for models and plots available at https://github.com/MPavFox/Socioecological-drivers-of-injuries/

## Author contribution

Conceptualization, M.A.P-F., L.J.N.B. and D.D.; Methodology, M.A.P-F., D.D., E.R.S. and S.E.; Resources, L.J.N.B., J.P.H., N.S-M., and A.R-L; Data Curation, M.A.P-F., C.M.K., N.R-B., J.E.N-D., and D.P.; Writing – Original Draft, M.A.P-F.; Writing – Review & Editing, M.A.P-F., D.D., E.R.S., L.J.N.B., C.M.K., J.P.H.; Supervision, D.D. and L.J.N.B.

