## Supplementary Information for "Socioecological drivers of injuries in female and male rhesus macaques (*Macaca mulatta*)"

**Supplemental Information**

### Supplementary Analyses

#### Group size and non-physical aggression between females

To test if group size influences the likelihood of females being non-physically aggressed (*i.e.,* submissive cues, threats) by other females in the group, we focused on female aggression data. Non-contact aggression data were excluded from the main analysis, but this data was part of the behavioural data collection protocol. In total, we collected 2285 events of non-physical aggression from a total of 422 females.

We specifically tested if the number of females in the group predicted the probability of a female receiving non-physical aggression from another female. Given that non-contact aggression encounters were common we used a Poisson distribution where the dependent variable was the count of non-contact aggression events received by a female in a given bimonthly interval. As independent variables, we included the number of females in the group, the reproductive season and an offset term for sampling effort (*i.e.,* hours of female focal followed). Random effects for animal ID and bimonthly intervals were also included.

We found that females living in larger groups were more likely to be non-physically aggressed by other females in the group (Fig. S4; Log-Odds fem count = 0.26, SE = 0.03, 89% CI = 0.21, 0.31; Table S10).

#### Sex ratio and non-physical aggression between females

To test if sex ratio influences the likelihood of females being non-physically aggressed (*i.e.,* submissive cues, threats) by other females in the group, we focused on female aggression data. Non-contact aggression data were excluded from the main analysis, but this data was part of the behavioural data collection protocol. In total, we collected 2285 events of non-physical aggression from a total of 422 females.

We specifically tested if the adult sex ratio (F:M) during the mating season predicted the probability of a female receiving non-physical aggression from another female. Given that non-contact aggression encounters were common we used a Poisson distribution where the dependent variable was the count of non-contact aggression events received by a female in a given bimonthly interval. As independent variables, we included the sex ratio and an offset term for sampling effort (*i.e.,* hours of female focal followed). Random effects for animal ID and bimonthly intervals were also included.

We found that females living in groups with female-biased sex ratios were more likely to be non-physically aggressed by other females in the group (Fig. S5; Log-Odds sex ratio = 0.11, SE = 0.05, 89% CI = 0.03, 0.18; Table S11).

### Supplementary Figures

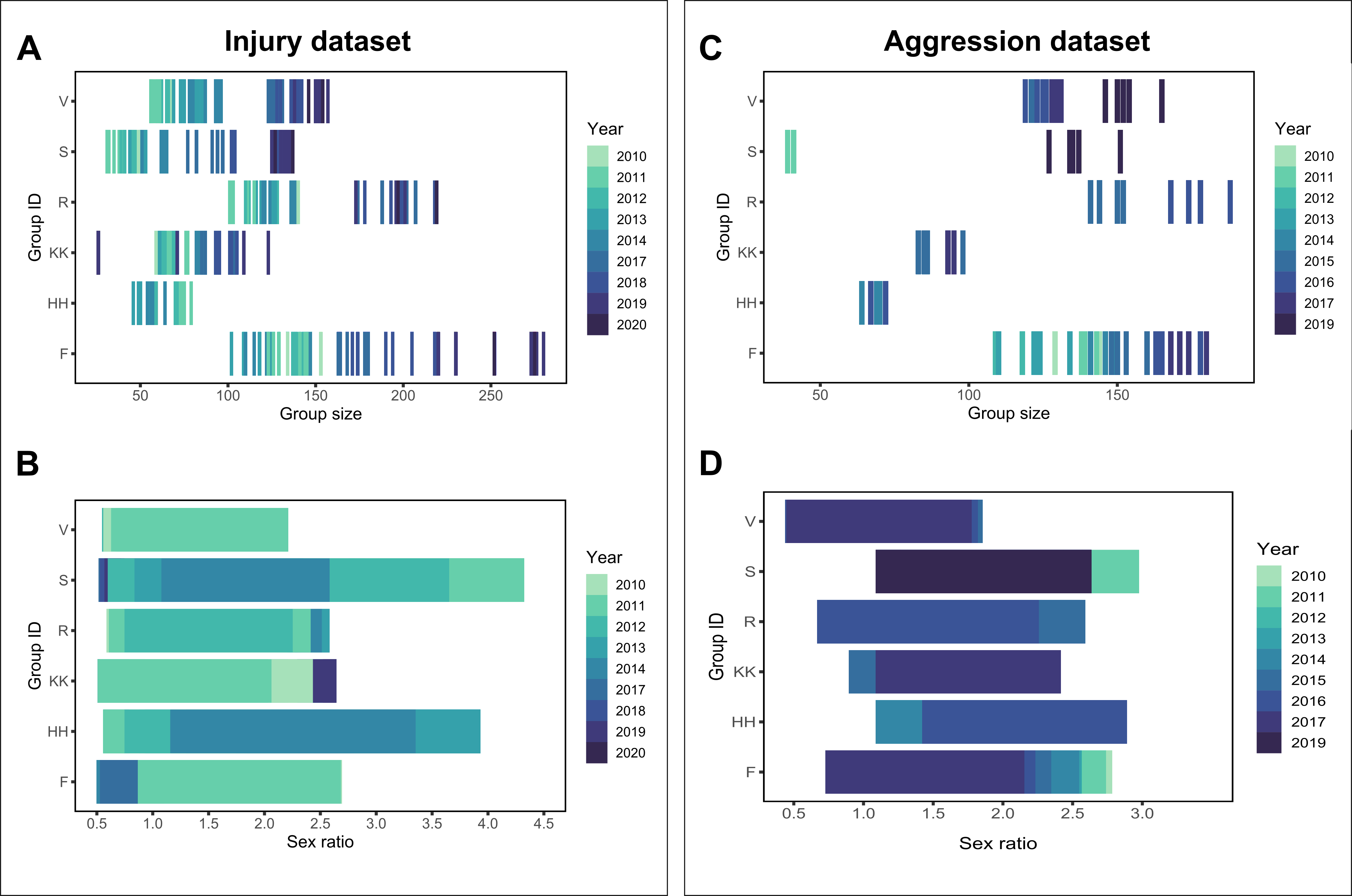

Figure S1: **Variation of group size and operational sex ratio per behavioural group across the 10 years of study.**

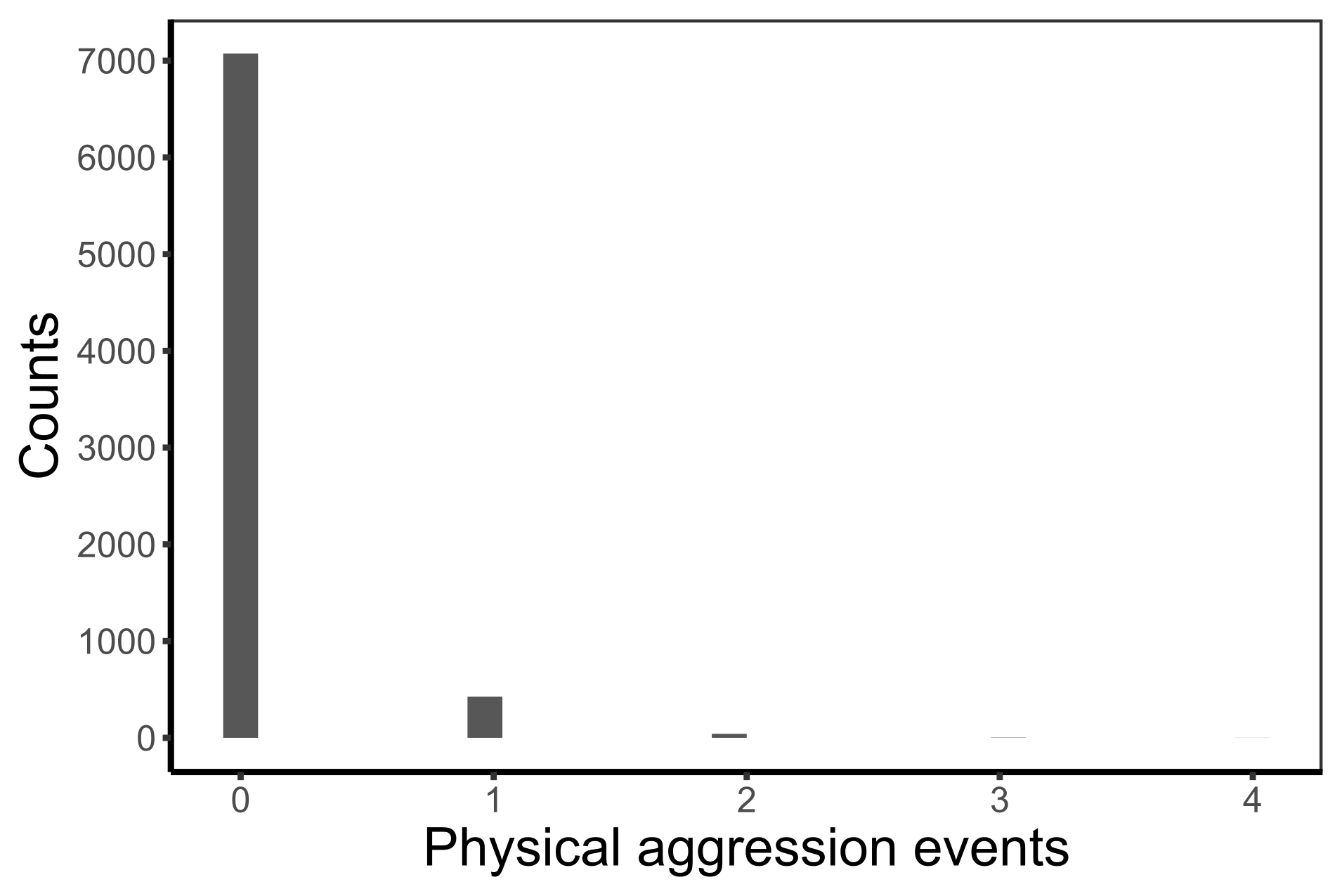

Figure S2: **Histogram of physical aggression received by a focal animal in a given bimonthly interval across the study period.**

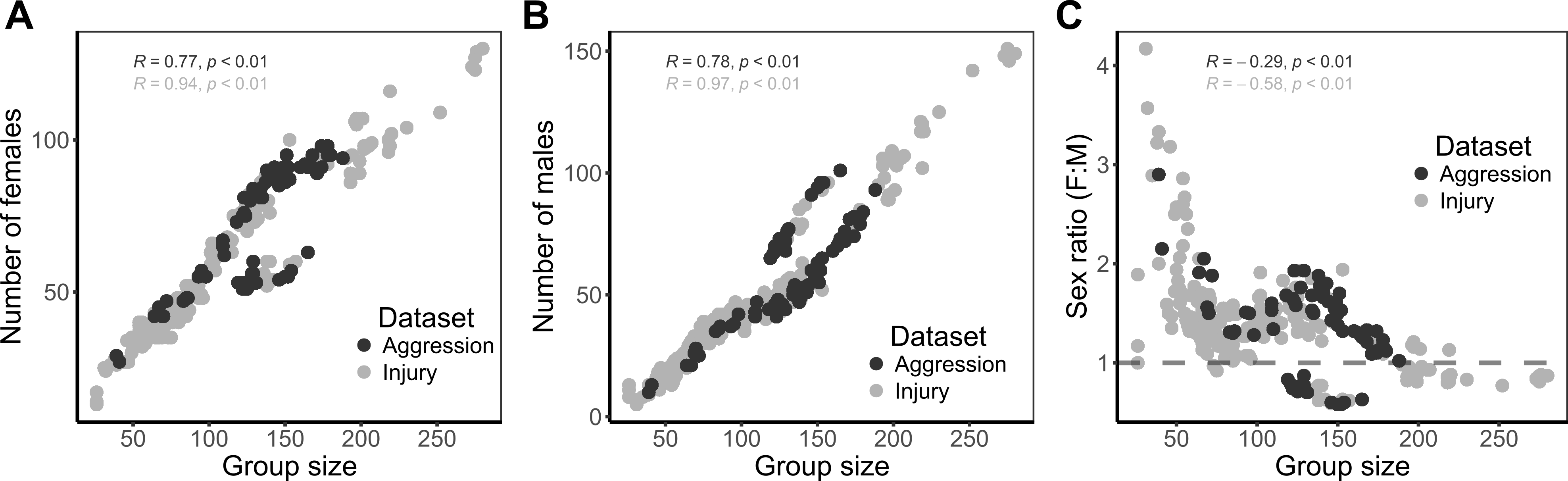

Figure S3: **Correlations of A) number of females, B) number of males and C) adult sex ratio with group size.**

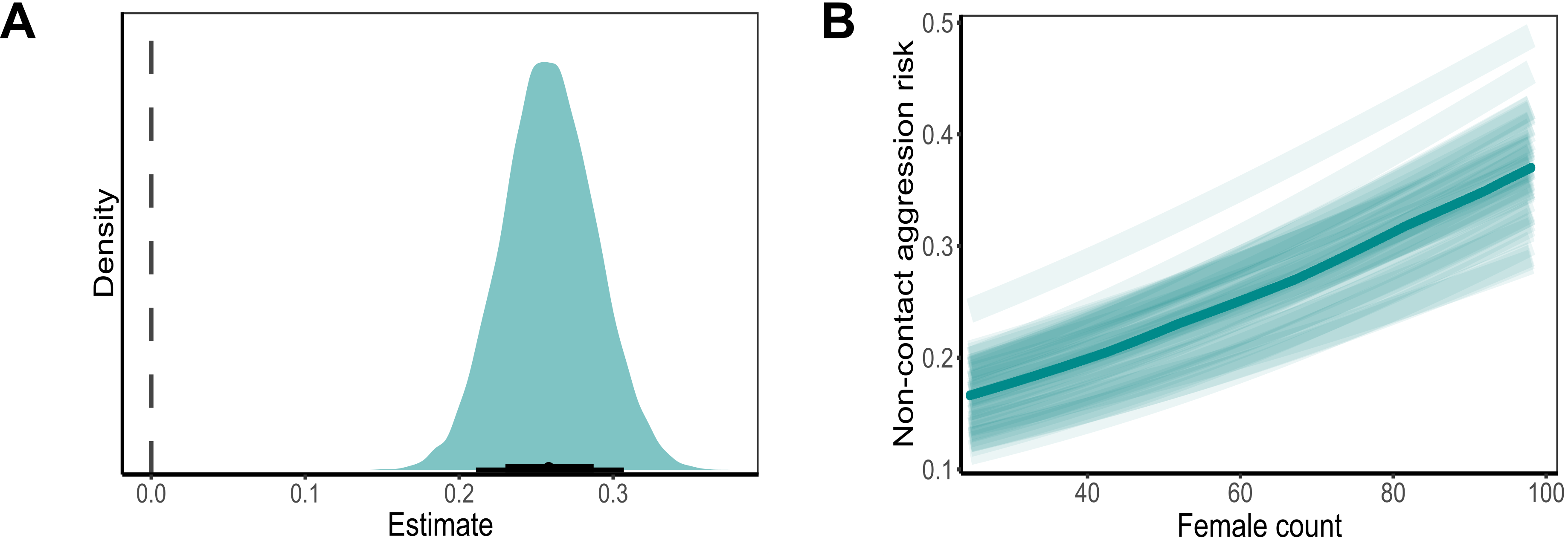

Figure S4: **Effect of the number of females on the risk of non-contact aggression from other females in the group. A)** Posterior distribution for the estimate of female count. Whisker indicates median, 89 % CI (thinner line) and 66% CI (thicker line). **B)** Predicted values for the risk of non-contact aggression from females as a function of the number of females in the group.

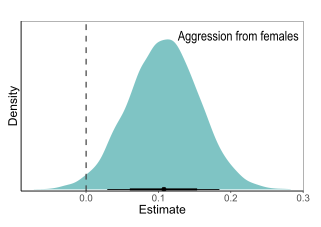

Figure S5: **Effect of sex ratio on a female’s risk of non-contact aggression from females in the group.**

**Table S1.** Output from logistic model predicting injury risk as a function of group size by sex.

|  | **Injury risk** | | |
| --- | --- | --- | --- |
| *Predictors* | *Log-Odds* | *std. Error* | *CI (89%)* |
| Intercept | -4.50 | 0.15 | -4.74 – -4.26 |
| scaled_group | 0.14 | 0.07 | 0.02 – 0.26 |
| sex: M | 0.29 | 0.08 | 0.16 – 0.43 |
| is_mating: is_mating1 | 1.22 | 0.22 | 0.86 – 1.58 |
| scaled_group:sexM | -0.36 | 0.08 | -0.49 – -0.23 |
| N _id_ | 1522 | | |
| N _year_bim_ | 46 | | |
| Observations | 28936 | | |

**scaled_group =** z-standardised group size, **sex : M** = males (females as intercept), **is_mating: is_mating1 =** reproductive season. Individual ID (**id**) and bimonthly interval (**year_bim**) were included as random effects. Model includes males and females.

**Table S2.** Output from logistic model predicting risk of female physical aggression as a function of the number of females in the group.

|  | **Risk of physical aggression** | | |
| --- | --- | --- | --- |
| *Predictors* | *Log-Odds* | *std. Error* | *CI (89%)* |
| Intercept | -4.08 | 0.18 | -4.39 – -3.82 |
| scaled_fem | -0.09 | 0.08 | -0.22 – 0.03 |
| is_mating: is_mating1 | 0.10 | 0.24 | -0.29 – 0.49 |
| N _id_ | 422 | | |
| N _year_bim_ | 39 | | |
| Observations | 4390 | | |

**scaled_fem =** z-standardised number of females in the group, **is_mating: is_mating1 =** reproductive season. Individual ID (**id**) and bimonthly interval (**year_bim**) were included as random effects. Model includes only females.

**Table S3.** Output from logistic model predicting risk of male physical aggression towards females as a function of group size.

|  | **Risk of physical aggression** | | |
| --- | --- | --- | --- |
| *Predictors* | *Log-Odds* | *std. Error* | *CI (89%)* |
| Intercept | -4.43 | 0.20 | -4.77 – -4.13 |
| scaled_group | -0.14 | 0.08 | -0.26 – -0.01 |
| is_mating: is_mating1 | 0.55 | 0.27 | 0.11 – 0.99 |
| N _id_ | 422 | | |
| N _year_bim_ | 39 | | |
| Observations | 4390 | | |

**scaled_group =** z-standardised group size, **is_mating: is_mating1 =** reproductive season. Individual ID (**id**) and bimonthly interval (**year_bim**) were included as random effects. Model includes only females.

**Table S4.** Output from logistic model predicting risk of male physical aggression as a function of the number of males in the group.

|  | **Risk of physical aggression** | | |
| --- | --- | --- | --- |
| *Predictors* | *Log-Odds* | *std. Error* | *CI (89%)* |
| Intercept | -4.22 | 0.23 | -4.63 – -3.88 |
| scaled_male | -0.06 | 0.13 | -0.27 – 0.15 |
| is_mating: is_mating1 | 0.68 | 0.30 | 0.20 – 1.18 |
| N _id_ | 326 | | |
| N _year_bim_ | 39 | | |
| Observations | 3154 | | |

**scaled_male =** z-standardised number of males in the group, **is_mating: is_mating1 =** reproductive season. Individual ID (**id**) and bimonthly interval (**year_bim**) were included as random effects. Model includes only males.

**Table S5.** Output from logistic model predicting risk of injury as a function of sex-ratio by sex during the mating season.

|  | **Injury risk** | | |
| --- | --- | --- | --- |
| *Predictors* | *Log-Odds* | *std. Error* | *CI (89%)* |
| Intercept | -3.27 | 0.15 | -3.53 – -3.01 |
| scaled_SR | -0.05 | 0.07 | -0.17 – 0.06 |
| sex: M | 0.54 | 0.10 | 0.38 – 0.70 |
| scaled_SR:sexM | 0.17 | 0.08 | 0.04 – 0.30 |
| N _id_ | 1419 | | |
| N _year_bim_ | 15 | | |
| Observations | 9286 | | |

**scaled_SR =** z-standardised sex-ratio (females to males in a group), **sex : M** = males (females as intercept). Individual ID (**id**) and bimonthly interval (**year_bim**) were included as random effects. Model includes males and females.

**Table S6.** Output from logistic model predicting risk of male physical aggression towards males as a function of sex-ratio during the mating season.

|  | **Risk of physical aggression** | | |
| --- | --- | --- | --- |
| *Predictors* | *Log-Odds* | *std. Error* | *CI (89%)* |
| Intercept | -4.35 | 0.29 | -4.93 – -3.94 |
| scaled_SR | 0.10 | 0.17 | -0.19 – 0.37 |
| N _id_ | 285 | | |
| N _year_bim_ | 13 | | |
| Observations | 1175 | | |

**scaled_SR =** z-standardised sex-ratio (females to males in a group). Individual ID (**id**) and bimonthly interval (**year_bim**) were included as random effects. Model includes only males.

**Table S7.** Output from logistic model predicting male aggression as a function of sex ratio considering only prime-age females (6-18) during the mating season.

|  | **Risk of physical aggression** | | |
| --- | --- | --- | --- |
| *Predictors* | *Log-Odds* | *std. Error* | *CI (89%)* |
| Intercept | -4.36 | 0.30 | -4.98 – -3.94 |
| scaled_SR2 | 0.04 | 0.19 | -0.27 – 0.34 |
| N _id_ | 285 | | |
| N _year_bim_ | 13 | | |
| Observations | 1175 | | |

**scaled_SR2 =** z-standardised sex ratio (prime-age females to males). Individual ID (**id**) and bimonthly interval (**year_bim**) were included as random effects. Model includes only males.

**Table S8.** Output from logistic model predicting risk of male physical aggression towards females as a function of sex-ratio during the mating season.

|  | **Risk of physical aggression** | | |
| --- | --- | --- | --- |
| *Predictors* | *Log-Odds* | *std. Error* | *CI (89%)* |
| Intercept | -3.81 | 0.21 | -4.22 – -3.52 |
| scaled_SR | -0.40 | 0.13 | -0.62 – -0.19 |
| N _id_ | 367 | | |
| N _year_bim_ | 13 | | |
| Observations | 1552 | | |

**scaled_SR =** z-standardised sex-ratio (females to males in a group). Individual ID (**id**) and bimonthly interval (**year_bim**) were included as random effects. Model includes only females.

**Table S9.** Output from logistic model predicting risk of female physical aggression towards females as a function of sex-ratio during the mating season.

|  | **Risk of physical aggression** | | |
| --- | --- | --- | --- |
| *Predictors* | *Log-Odds* | *std. Error* | *CI (89%)* |
| Intercept | -4.12 | 0.30 | -4.69 – -3.68 |
| scaled_SR | 0.02 | 0.15 | -0.22 – 0.27 |
| N _id_ | 367 | | |
| N _year_bim_ | 13 | | |
| Observations | 1552 | | |

**scaled_SR =** z-standardised sex-ratio (females to males in a group). Individual ID (**id**) and bimonthly interval (**year_bim**) were included as random effects. Model includes only females.

**Table S10.** Output from Poisson model predicting risk of female non-contact aggression towards females as a function of the number of females in the group.

|  | **Risk of non-contact aggression** | | |
| --- | --- | --- | --- |
| *Predictors* | *Log-Mean* | *std. Error* | *CI (89%)* |
| Intercept | -0.87 | 0.16 | -1.13 – -0.61 |
| scaled_fem | 0.26 | 0.03 | 0.21 – 0.31 |
| is_mating: is_mating1 | 0.14 | 0.26 | -0.30 – 0.57 |
| N _id_ | 422 | | |
| N _year_bim_ | 39 | | |
| Observations | 4390 | | |

**scaled_fem =** z-standardised number of females in the group, **is_mating: is_mating1 =** reproductive season. Individual ID (**id**) and bimonthly interval (**year_bim**) were included as random effects. Model includes only females.

**Table S11.** Output from logistic model predicting the risk of female non-contact aggression towards females as a function of sex-ratio during the mating season.

|  | **Risk of non-contact aggression** | | |
| --- | --- | --- | --- |
| *Predictors* | *Log-Mean* | *std. Error* | *CI (89%)* |
| Intercept | -0.81 | 0.20 | -1.16 – -0.47 |
| scaled_SR | 0.11 | 0.05 | 0.03 – 0.18 |
| N _id_ | 367 | | |
| N _year_bim_ | 13 | | |
| Observations | 1552 | | |

**scaled_SR =** z-standardised sex-ratio (females to males in a group). Individual ID (**id**) and bimonthly interval (**year_bim**) were included as random effects. Model includes only females.
